# A Note on Stochastic Modeling of Biological Systems: Automatic Generation of an Optimized Gillepsie Algorithm

**DOI:** 10.1101/395392

**Authors:** Quentin Vanhaelen

## Abstract

Signaling pathways and gene regulatory networks (GRNs) play a central role in the signal transduction and regulation of biochemical processes occurring within the cellular environment. Understanding their mechanisms and dynamics is of major interest in various areas of life sciences and biological sciences. For example controlling stem cell fate decision requires a comprehension of the dynamical behavior of the networks involved in stem cell differentiation and pluripotency maintenance. In addition to analytical mathematical methods which are applicable for small or medium sized systems, there are many computational approaches to model and analyze the behavior of larger systems. However, from a dynamical point of view, modeling a combination of signaling pathways and GRNs present several challenges. Indeed, in addition to being of large dimensionality, these systems have specific dynamical features. Among the most commonly encountered is that the signal transduction controlled by the signaling pathways occurs at a different time scale than the transcription and translation processes. Also, stochasticity is known to strongly impact the regulation of gene expression. In this paper, we describe a simple implementation of an optimized version of the Gillespie algorithm for simulating relatively large biological networks which include delayed reactions. The implementation presented herein comes with a script for automatically generating the different data structures and source files of the algorithm using standardized input files.

**Code availability:** The Fortran90 implementation of the code and the R script described here as well as the tutorial with practical instructions are stored on the following github repository qvhaelen/ typhon

## 1 Introduction

Using the new proteomic [1] and genomic data available, it is possible to study the organization of the cellular environment and its components [2, 3]. A current challenge concerns the elaboration of a comprehensive view of how these different components interact together. This includes the understanding of protein-protein interactions and gene regulation taken separately as well as the interactions between the proteosome and the transcriptome. This is not a trivial task because proteomic and transcriptomic data are obtained by different well defined experimental protocols. Studies are undertaken to establish dynamical links between these two types of data and although many questions remain unanswered, recent encouraging results have been obtained for example about the relationships between mRNA and protein concentrations [4, 5, 6, 7]. It is well-known that considering the complexity of the interactions taking place within the cellular environment, gaining an accurate understanding of the cellular biochemical processes requires a systemic approach where the cell is considered as a complex dynamical system [8]. Within this framework, it is possible to build a systemic description of the biological processes in terms of their functional and dynamical properties [9, 10]. From a practical point of view, a systemic approach is also necessary to understand how a local dysfunction of a small set of molecules affecting a restricted number of defined epigenetic [11] and metabolic processes [12, 13] may propagate to all parts of the cell leading to a progressive disruption of the general homeostasis [14]. The studies performed so far provide an interesting picture of how the cellular components are organized as dynamical motifs and cycles [15, 16]. These dynamical motifs communicate together to achieve specific tasks. Inside the nucleus for example, the transcription factors interact together forming structured dynamical patterns called gene regulatory networks. These networks control and modulate genes expression via promoter silencing but supplementary mechanisms such as mRNA splicing [17, 18], chromatin remodeling [19] and epigenetic changes also intervene. Epigenetic modifications are a set of elaborate dynamical adaptations of the structure of the chromatin [20, 21] which contribute significantly to the regulation of gene transcription [22, 23, 24, 25].

Understanding and controlling the interaction between external perturbation, signal transduction and biological response is especially important within the field of stem cell research which focuses on understanding the two main mechanisms forming the backbone of stem cell fate decision, i.e., Cellular differentiation and pluripotency maintenance. Observations show that pluripotency state maintenance is a function of the external environment and input received. A dynamical balance between environmental clues and cell signals is required to preserve the self-renewal and tissue regenerative capacity of stem cells. Stem cell commitment and differentiation into a specific cell lineage is induced by modifying this dynamical balance with new external perturbations in order to activate or inhibit specific sets of signaling pathways which transmit external inputs to the core of trasnscription factor (TF) networks. Indeed, it is now experimentally established that pluripotency maintenance is essentially under the control of transcription factors [26, 27, 28] which act as master regulators of highly connected transcription networks [29, 30, 31, 32]. These TFs are continually attempting to specify differentiation to their own lineage of interest. This is the main reason why direct external intervention, through activation or inhibition of one or several signaling pathways is required to reinforce the pluripotent state or to drive differentiation [33, 34, 35]. Thus, the pluripotency state could be considered as a metastable state. The maintenance of the pluripotency state being a function of the external environment, any realistic dynamical description should include the cellular components responsible for processing this external signal. The TF core network being localized inside the nucleus, it receives this external signal through a second network of signaling pathways located inside the cytoplasmic compartment. Thus, it is essential to integrate The TF network and the signaling pathways within a single framework [36]. The behavior of the resulting extended regulatory networks can be better understood when defined as a complex dynamical system [8].

This discussion illustrates the importance of being able to simulate the dynamics of relatively large systems including those eliciting properties such as stochasticity and multiple scale dynamics. Although other computational methods such as constraint-based methods or pathway perturbation analysis methods using pathway maps can handle models of large dimensionality, kinetic-based models are the most appropriate for a detailed study of the dynamical features of the model [37, 38, 39]. Many works have been published regarding stochastic kinetic modeling of biological systems. Reviewing those contributions is outside the scope of this paper. The aim of this work is to shortly describe a stand alone implementation of an optimized version of the exact Gillespie algorithm. This code comes with a practical methodology to describe simple kinetic models which can be used as standardized inputs by a specifically designed R script to automatically generate the source files needed to perform the simulation. The complete description of the R script and a template of the standardized input files are stored with the code. In what follows, the mathematical framework to build the kinetic models is shortly described with a emphasis on the types of reactions allowed. Then, the algorithmic method and the mean features of its implementation are summarized. This is followed by the discussion of a small case study using a single signaling pathway as an example.

## 2 Definition of the system

### 2.1 Mathematical formulation

An extended regulatory network is a dynamical system of biochemical components interacting together through a set of biochemical reactions. In order to allow the easy combination of various pathways together in a flexible way, the kinetic is described using a scheme based on the law of mass action. It offers a good mathematical stability and easy systematic interpretation in terms of biological processes. Considering the nature of the dynamics encountered in signaling cascades and gene regulatory networks, all reactions are described in terms of elementary processes and other more specific kinetic schemes and approximations will not be considered.

Formally, when analyzing the dynamics of a biological network, one is concerned with the temporal evolution of the set of *m* species^1^ *X*_*i*_ (*i* = 1, *…, m*) which composes the network. These species react together through a set of *r* chemical elementary reactions. The derivation of the dynamics of the network is based on the law of mass action which states that, for an elementary reaction, that is, a reaction in which all of the stoichiometric coefficients of the reactants are one, the rate of reaction is proportional to the product of the concentrations of the reactants [40]. Although the mass action kinetics scheme expands the dimension of the model, it provides a regularized and simplified mathematical structure. This feature makes further analysis more tractable. Formally, one defines the stoichiometric matrix 𝒩_*ij*_ = *p*_*ij*_ *- r*_*ij*_ with (*i* = 1, *…, m*) and (*j* = 1, *…, r*) and the reaction rate vector as 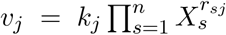 with (*j* = 1, *…, m*) where the *p*_*ij*_ term is the stoichiometric coefficient for the *i-*th species in the *j-*th reaction if appearing on the right of the reaction arrows, *r*_*ij*_ if on the left; *k*_*j*_ is the forward rate constant [41]. Using 𝒩 and 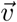 the function 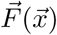 is built such that 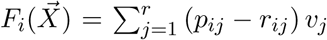 (i = 1, …, *m*). Each 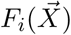 represents the contribution of the reactions acting on the species *X*_*i*_.

In our description all the chemical reactions occurring in the extended regulatory network are modeled using the following elementary reactions^2^. Association (between two different molecules or between similar molecules (dimerization)) *X*_1_ + *X*_2_ *→ X*_3_, dissociation (giving two identical molecules or two similar molecules) *X*_3_ *→ X*_1_ + *X*_2_, creation (this reaction mimic the dynamic of molecules for which no explicit creation pathway is included) ∅ *→ X*, degradation (last step of any degradation process including lysosomal and proteosomal pathway) *X→*∅, direct activation/inhibition (when enzyme can be neglected, like phosphatase, etc.) *X→X*^*\**^. Nevertheless, For any chemical process involving enzymes (E) which can not be ignored we use the Michaelis-Menten scheme:

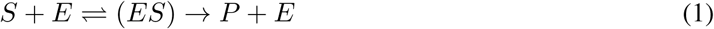

Where *S* is the substrate and *P* the product of the reaction. We model these three reactions independently as follows:

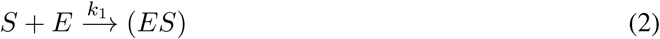

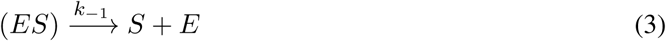

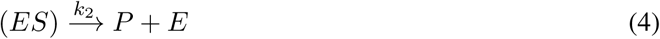

The parameters appearing in the Michaelis-Menten formula

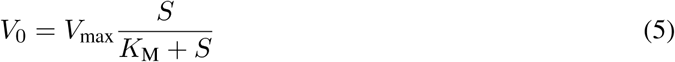

are connected to the rate constants as follows

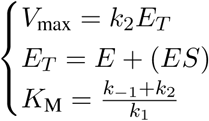

In general, the reversible chemical reactions are always modeled as a set of separate irreversible forward reactions. The set of reactions described above is enough to describe any kind of process occurring in most of the signaling pathways.

The dynamics of GRNs is mainly concerned with the regulation of gene expression [42, 43, 44, 45, 46]. In this work, DNA and ribosome are not explicitly taken into account in the kinetic equations. Ribosome could be included with a complete description of ribosomal proteins assembly but, as discussed below, it leads to much more complexity. We propose to model the regulation of gene expression using the following scheme:

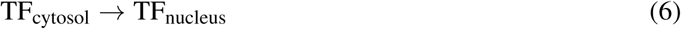

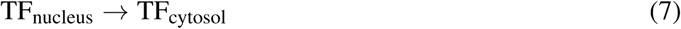

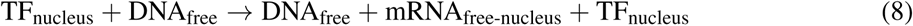

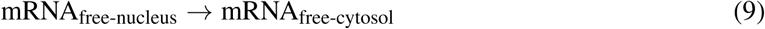

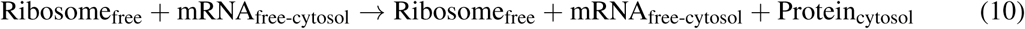

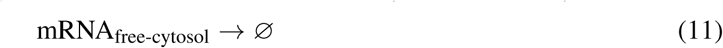

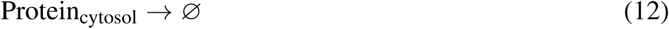

When a gene is repressed by one or several genes binding the promoter site, the effect can be included using a modification of the transcription rate constant as follows:

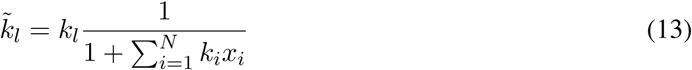

Where *k*_*l*_ is the non modified transcription rate, *N* the number of genes that can repress the promoters, *x*_*i*_ the concentration in the nucleus of repressor *i* and *k*_*i*_ the intensity of the repression.

### 2.2 GRNs and stochasticity

Experimental evidences show that a lot of mechanisms occurring in a cell and especially within GRN elicit stochastic properties. For example, stochastic expression of key genes (Nanog, Sox2, Oct4,) can lead to heterogeneous populations of stem cells even when starting from a homogeneous culture in a homogeneous environment. In a regulatory network, stochasticity can have different origins including the low number of molecules usually involved in molecular processes occurring in a cell, the effect of external perturbations and response of signaling pathways. Furthermore, the transcription/translation processes are also stochastic with an impact on the release of proteins. It has also been shown that dynamical sub-networks can increase (positive feedback) or reduce (negative feedback) stochastic effects. Taking into account the stochastic nature of the dynamics is thus of primary importance when simulating the dynamics of an extended regulatory network and appropriate methods must be chosen.

### 2.3 Extended regulatory network and multi-scale processes

Molecular events, such as dimerization and phosphorylation, occurring in signaling pathways and processes such as TF binding, transcription/ translation and mRNA transfer occur on different timescales. Regarding the transcription and translation, one observes a delay between the beginning of these processes and the release of the final product. It is known that systems involving delayed dynamics can elicit specific behavior [47, 48]. From a computational perspective, including dynamical events occurring at various timescale within a single dynamical model can be complex. Two approaches can be used. If one stands from the point of view of the slowest processes, one will assume that the fastest processes have already reached their equilibrium state when the slowest events take place. Otherwise, one takes the point of view of the fastest processes and in that case, one needs to take into account the fact that there is a time delay between the release of the products by the fastest reactions and the release of the product by the slowest reactions. It is this second point of view which is considered in this work. Thus, we divide the set of reactions into two distinct subsets. Firstly, we assume that the fastest chemical reactions occurs immediately (products are released at once). Secondly, the slower chemical reactions are supposed to release their products with a delay which must be specified. Thus we need to know the delay between the beginning and the release of the products for the slower processes. The exact duration of transcription rate and translation rate usually varies from one gene to another and other chemical or physiological parameters can intervene. Nevertheless, these rates are always slower than the dimerization mechanisms. Modeling multi-scale stochastic systems can be done using dSSA (exact formulation of a SSA including delay, see description below) which allows considering the delay when the model includes processes occurring on different time scales.

## 3 Stochastic simulation: an optimized version of the Gillespie algorithm

There are two formalisms for mathematically describing the time behavior of a spatially homogeneous chemical system. The deterministic approach regards the time evolution as a continuous, wholly predictable process which is governed by a set of coupled ODEs. The stochastic approach regards the time evolution as a kind of random-walk process which is governed by a single differential-difference equation (the master equation). In practice, many modeling studies rely on using different flavors of ODE-based formulations, probably because many scientists are more familiar with purely deterministic approaches and that formulating a model in terms of ODEs and solving this system is most of the time straightforward. Although, deterministic modeling relies on several strong assumptions, it can provide suitable and accurate results in many situations. Furthermore, formulating the dynamics of the system in terms of ODEs makes easier to perform additional analytical or numerical analysis of the underlying dynamical properties of the system.

Moreover, when considering dynamical systems eliciting stochastic behavior, such as GRNs or signaling pathways whose dynamics is directly connected to a GRN, fairly simple kinetic theory arguments show that the stochastic formulation of chemical kinetics has a firmer physical basis than the deterministic formulation. However, the fundamental stochastic master equation becomes quickly mathematically and computationally intractable when the dimensionality of the system becomes large, i.e., for systems with more than 5 or 6 variables. This problem is well known and different approaches have been suggested to overcome this issue [49, 50, 51, 52, 53, 54]. One of the most famous is the Gillespie method, also known as the stochastic simulation algorithm (SSA), which allows making exact numerical calculations within the framework of the stochastic formulation without having to deal with the master equation directly [55]. It uses a rigorously derived Monte Carlo procedure to simulate the time evolution of the given chemical system. Like the master equation, the SSA correctly accounts for the inherent fluctuations and correlations that are necessarily ignored in the deterministic formulation. In addition, unlike most procedures for numerically solving the ODE, this algorithm never approximates infinitesimal time increments.

The SSA or Gillespie algorithm, were initially described in the fundamental paper [55]. The key steps of this algorithm, called the direct method (DM), can be summarized as follows:

**Step** 0: (Initialization). Input the desired values for the *M* reaction constants *c*_1_, *…, c*_*M*_ and the *N* initial molecular population numbers *X*_1_, *…, X*_*N*_. Set the time variable *t* and the reaction counter *n* both to zero. Initialize the Pseudo Random Number Generator (PRNG).

**Step** 1: Calculate and store the *M* quantities *a*_*µ*_ = *c*_*µ*_*h*_*µ*_, (where *h*_*µ*_ is the number of distinct molecular reactant combinations available in the state for reaction *µ*), for the current molecular population numbers. Also calculate and store as *a*_0_ the sum of the *M a*_*µ*_ values.

**Step** 2: Generate two random numbers *r*_1_ and *r*_2_ using the PRNG, and calculate *τ* and *µ* according to

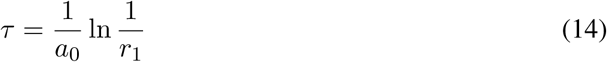

and

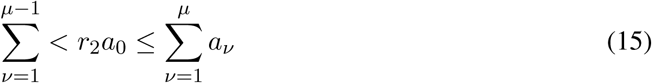

**Step** 3: Using the *τ* and *µ* values obtained in step two, increase *t* by *τ*, and adjust the molecular population levels to reflect the occurrence of one *R*_*µ*_ reaction. Then increase the reaction counter *n* by 1 and return to step one.

The SSA is a very accurate method but when the number of reactions becomes large, the DM becomes too slow to be useful in practice. It is necessary to use several specific algorithmic techniques which can speed up the algorithm while keeping its accuracy intact. Here we provide a general description of the different approaches used in our implementation. The strategy is to focus on the steps of the DM which are the most computationally expensive.

**Dependency graph** In the DM, after a reaction has been fired, all the *a*_*µ*_ coefficients are updated. It easy to see that only a few of them are actually modified when a reaction occurs. An appropriate way to take this fact into account is to implement a dependency graph. For each reaction, this data structure stores the label of the *a*_*µ*_ which are modified. For each reaction, the set of *a*_*µ*_ to be updated is constructed as follow: we consider the label of the species whose belongs to the set of reactants or products of the given reaction. If any of these species appears in the set of reactants of a reaction *i*, then the corresponding *a*_(*µ*=*i*)_ is added in the set of *a*_*µ*_ to be updated. It is also useful to save the smallest label which must be updated [56].

**Partial Summation** Another expensive step in the DM is the computation of *a*_0_. It is possible to reduce the number of operations by building a set of partial sum over the *a*_*µ*_. For a system with *M* chemical reactions, a set of *M* partial sums is constructed. A partial sum is defined as follows:

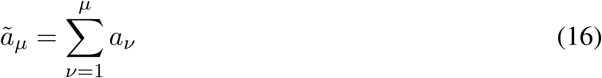

Using a recursive formula it is possible to compute each partial sum from the previous one. Taking into account that 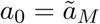, and the fact that we know the smallest label 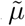 which has been modified it is possible to restrict the computation of *a*_0_ to the subset of partial sums 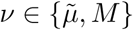.

**Binary tree data structure and binary tree search** Locating the reaction to fire next is the most expensive step of the DM. A way to reduce the cost of this step is to make use of a specific data structure called binary tree and the related binary tree search algorithm [57]. This approach is known as the Logarithmic Direct Method (LDM) [58] and the use of binary tree search reduces the search depth to *O*(log*M*). Furthermore, for LDM, the average search depth is independent of the ordering of the reactions. Hence there is no need for a pre-simulation.

**Contracting the stoichiometric matrix** Another step of the DM which is computationally expensive is the update of the number of molecules of each species. This update depends on the label of the reactions fired. In the direct method the stoichiometric coefficients of the chemical reactions are stored in a matrix whose dimensions scale with the number of reactions and species of the system. A key property of this matrix is that it is usually sparse, that is, for each reaction there are only a few species which must be updated. It means that in the DM, most of the operations are additions of zero terms. A way to reduce the computational cost of the update is to use sparse matrix techniques. In our case we have implemented a supplementary structure which contains for each reaction the label of the non-zero stoichiometric coefficients. Thus the update is done by taking into account the labels appearing in this structure.

In addition to optimizing the performances of the standard SSA, supplementary adaptations must be done to include the effects of multi scale dynamics. Indeed, in the initial formulation of the Gillespie method, it is assumed that all reactions occur instantly, i.e., products are released instantly. While this is true in many cases, it is also possible that some chemical reactions take certain time to finish after they are initiated. Thus, the product of such reactions will emerge after certain delays. It is important to distinguish two different kinds of delayed reactions. The *non consuming reaction*, where the reactants of an unfinished reaction can participate in a new reaction and the *consuming reaction*, where the reactants of an unfinished reaction cannot participate in a new reaction.

The main difference between consuming and non consuming reactions occurs at the level of the evolution of the reactant: When a non consuming reaction occurs, the population of the reactants does not change and the involved species can participate in a new reaction; however, when a consuming reaction occurs, the population of the reactants changes immediately, and the molecules involved in a consuming reaction can not participate in a new reaction.

There are many algorithms which have been released to integrate delayed reactions into the standard SSA scheme. However, many of them rely on various approximations which although allowing to mimic the effects of delayed reactions, do not constitute a mathematically accurate implementation of delayed reactions with th SSA. In this work, the algorithm chosen [59], is an exact SSA algorithm for chemical reaction systems with delays. The algorithm is exact, since it is rigorously developed based upon the fundamental premise of stochastic chemical kinetics derived by Gillespie. The difference between the complexity of the algorithm [59] and other exact methods like rejection algorithm mainly lies in the number of random variables generated by two algorithms. Specifically, the rejection algorithm needs to generate up to 50% more random variables than the algorithm presented in [59], if the chemical reaction system is dominated by reactions with delays. On the other hand, if the reaction system is dominated by reactions without delays, the rejection algorithm still generates slightly more random variables than the algorithm [59].

## 4 Examples of simulation

### 4.1 The model: The LIF pathway

The model is based on a work published by Mahdavi et al. [60] to characterize the dynamics of the LIFR-GP130 signaling pathway. This pathway is well known as a key regulator involved in the pluripotency maintenance of mouse embryonic stem cell [44]. Initially formulated as a set of ODEs, this model includes the formation of the activated receptor complex between LIFR and GP130 as well as its trafficking along the membrane and the effects of the negative feedback loop by SOCS3. The second part of the model is the activation of the STAT3 dimer via endogenous and exogenous phosphorylation. Activation of STAT3 dimer is followed by its nuclear translocation where it acts as a transcription factor for SOCS3 and other key pluripotency factors such as NANOG and OCT4. Moreover, it has been recently shown that NANOG is also involved in a positive feedback loop to increase STAT3 signaling by inhibiting SOCS3 transcription [61]. Although rather simple, this model is useful as it provides the user with a concrete example of how the different types of reactions described above should be implemented to use the code described herein. In what follows, we use the results of simulations performed with the SSA to illustrate the main dynamical features of this model.

### 4.2 Dynamics without multi-scale dynamics

The simulation starts without ligand and only GP130 receptors, LIF receptors and cytoplasmic STAT3 dimers are present. The model assumes that endogenous phosphorylation of cytoplasmic STAT3 dimers (Tyr705-phosphorylated 2STAT3) takes place. Thus as chemical reactions occur, the concentration of phosphorylated STA3 dimers (p2STAT3) increases inside the cytoplasm and can translocate inside the nucleus. Although present at low concentration, the level of p2STAT3 inside the nucleus is enough to induce the basal transcription of SOCS3 mRNA. SOCS3 mRNA can translocate into the cytoplasm where translation of SOCS3 protein occurs. Once the system has reached its equilibrium, LIF inducer is added. From a dynamical point of view, addition of LIF can be considered as an external perturbation which is applied on the cellular network^3^. LIF first binds to LIFR receptor which then form a receptor complex with GP130 co-receptor. Figure 1 shows the decrease of LIFR and GP130 concentration upon addition of external ligands for various concentrations.

**Figure 1:**
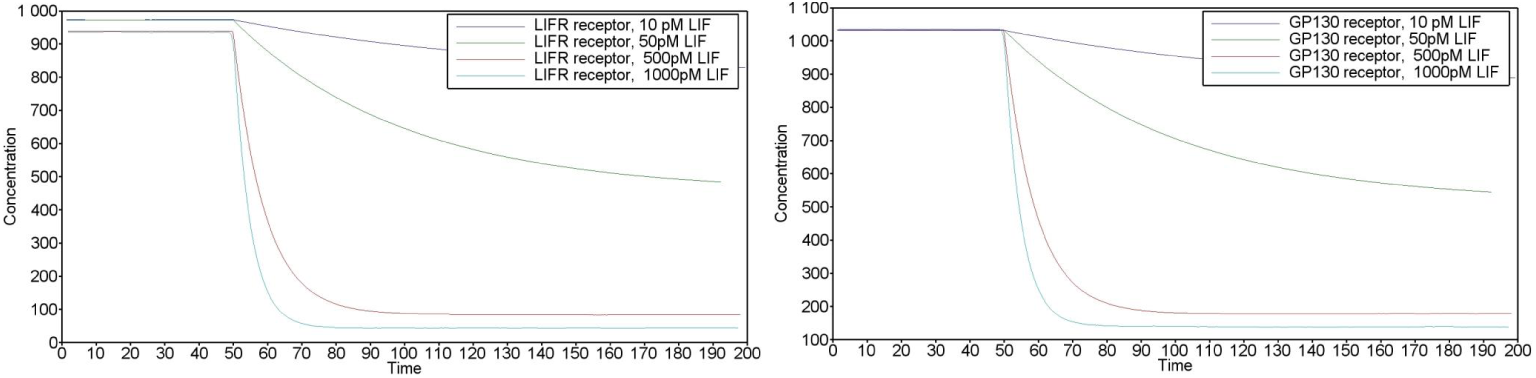
Response of receptors with respect to addition of LIF inducer. Response of LIFR (Left). Response of GP130 (Right).

Once the formation and activation of the LIFR-GP130-JAK is achieved, STAT3 dimers bind to the complex and become phosphorylated. As illustrated on Figure 2, the equilibrium switches from a state of a high concentration of cytoplasmic STAT3 dimer combined with a low concentration of nuclear p2STAT3 to a new state characterized by a lower concentration of cytoplasmic 2STAT3 with a higher concentration of nuclear p2STAT3.

**Figure 2:**
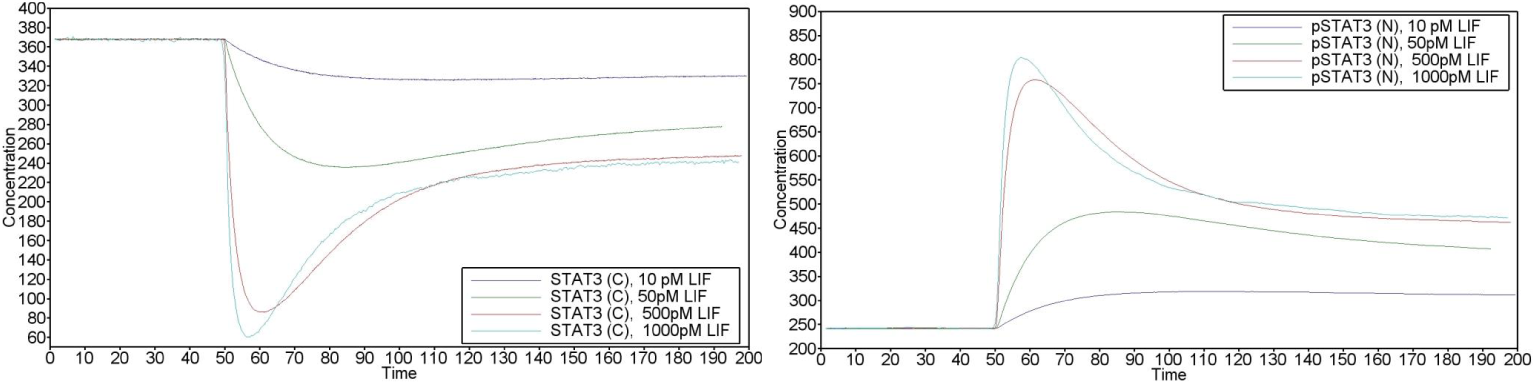
Evolution of the 2STAT3 concentration upon LIF addition in the cytoplasm (Left). LIF addition leads to an increase of the activated form of 2STAT3 dimer concentration inside the nucleus (Right)

Higher concentration of p2STAT3 inside the nucleus leads to a significant increase of SOCS3 mRNA, see Figure 3. As a result, level of cytoplasmic SOCS3 increases. Upon addition of LIF, the negative feedback loop established between SOCS3 proteins and the receptor complex plays an important in the regulation of the signal strength. This is essentially because binding of cytoplasmic SOCS3 to the LIFR-GP130-JAK complex is the main step leading to the final degradation of the receptor complex. The strong decrease in SOCS3 mRNA concentration which is observed right after LIF addition is the direct consequence of the activation of this feedback, Fig. 3.

**Figure 3:**
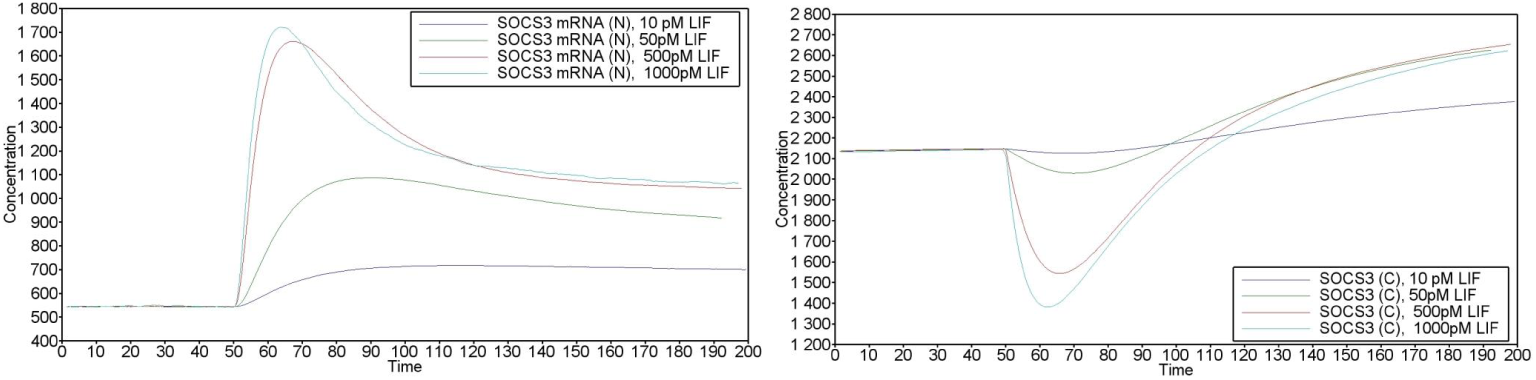
Following LIF addition, transcription of SOCS3 increases. SOCS3 mRNA in the nucleus (Left). SOCS3 protein concentration inside the cytoplasm (Right)

The purpose of the negative feedback loop is to improve the stability of the system by reinforcing the regulation of the signal. In the situation analyzed here, the feedback loop allows the degradation of the receptor complex which in turn induces the reduction of the exogenous phosphorylation of STAT3 dimers. As a result, nuclear p2STAT3 concentration decreases from its highest level almost directly after LIF addition, Fig. 2, and the system reaches a new equilibrium state characterized by higher level of p2STAT3 and SOCS3 (C). The kinetic parameters controlling the rate of association and dissociation of the receptor complexes as well as the translocation rate of 2STAT3 have a critical influence on the behavior of the cellular network.

The model presented here also contains the PIAS3 inhibitor. PIAS3 is present inside the cytoplasmic environment as well as inside the nuclear compartment. In the cytoplasmic compartment, PIAS3 acts as an inhibitor by forming a complex with the activated form of 2STAT3 preventing its translocation into the nucleus. In the nucleus, PIAS3 may also binds to the 2STAT3 preventing its export and further activation by phosphorylation. As a consequence, the level of available STAT3 within the phosphorylation loop is reduced. This second mechanism also contributes to the regulation of the signal strength.

Upon withdrawal of LIF, the ability of the cellular network to come to back to its initial state is a function of the kinetic parameters controlling the deactivation and degradation of the receptor complexes. Thus the frequency of addition and withdrawal of LIF can have an impact on the level of cytoplasmic p2STAT3 and SOCS3 protein. To illustrate this, we show the behavior of the main components of the cellular network for a periodical addition and withdrawal of LIF on Fig. 4.

**Figure 4:**
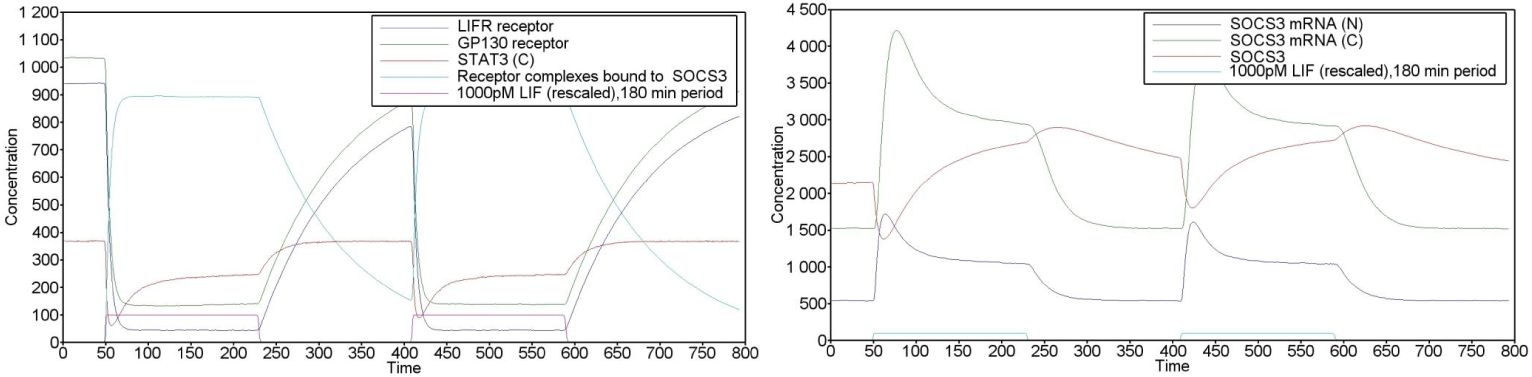
Temporal evolution of the main components of the pathway for a periodical addition of LIF inducer. Data are shown for a periodical addition of 180 minutes.

### 4.3 Multi-scale dynamics: effects of delayed reactions

To conclude we illustrate the effects of adding delay within the dynamical system for some reactions. We consider a delay for the transcription of SOCS3-mRNA (1.58 min), for the nuclear export of SOCS3-mRNA (1.92 min), for the import and export of the STAT3 dimer (1.2 min and 1.4 min) and for the translation of SOCS3 protein (100 min). The results are shown of Fig. 5 and Fig. 6. In order to understand how the imposed delay can affect the dynamics, the results are shown for different values of the delay for the translation of SOCS3. As expected, the addition of delay impacts how the system react to the LIF addition but also the concentration levels corresponding to the stationary states with and without LIF. The saturation observed for the activated form of the STAT3 dimer (Fig. 5 bottom) and SOCS3-mRNA inside the cytoplasm (Fig. 6 top) and the complete absence of SOCS3 proteins (Fig. 6 bottom) upon LIF addition varies according to the value chosen for the delay of the SOCS3 translation. From a dynamical point of view thecounter-reaction of the system to the absence of SOCS3 protein due to very large delay is a direct effect of the negative feedback loop involving SOCS3 described above. Indeed, upon LIF addition, the complex formed by LIFR and GP130 can be activated and trigger, without delay, the degradation of the amount SOCS3 already present within the system. As before, there is also a translocation of a higher concentration of activated STAT3 dimer which can increase SOCS3-mRNA transcription. However, because the time required to release SOCS3 (up to 100 minutes) is significant, the increase in SOCS3 transcription cannot directly compensate for the degradation immediately taking place. When all SOCS3 is eliminated from the system, this feedback loop ceases to function. This switches the equilibrium for the concentration levels of activated STAT3. When SOCS3 proteins begin to be released, the system undergoes a dynamics similar to the case without delay with a strong increase of SOCS3 proteins and a strong decrease of activated STAT3 dimer, the behavior being more accentuated as the value of the delay increases. The system finally reaches a new steady state which takes longer to be established than in the case without delay.

**Figure 5:**
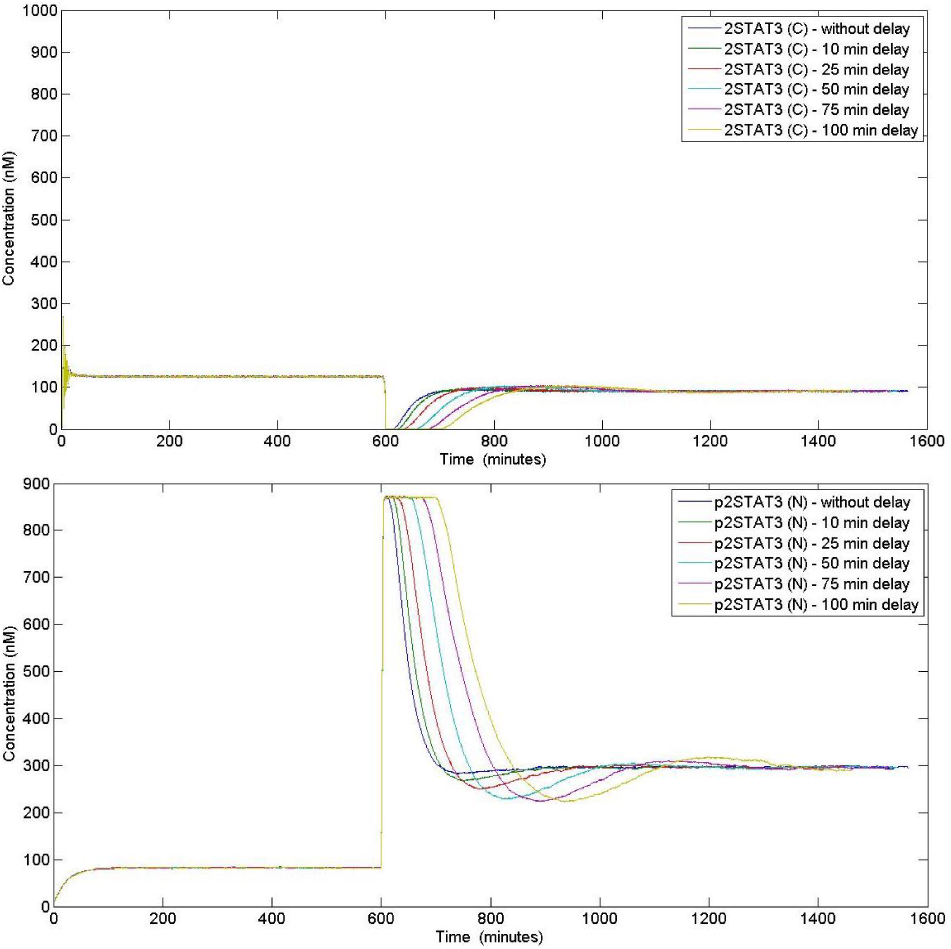
Temporal evolution of the dimer STAT3 inside the cytoplasm (Up) and for the activated dimer STAT3 inside the nucleus (Bottom) for different values of the delay for the translation of SOCS3.

**Figure 6:**
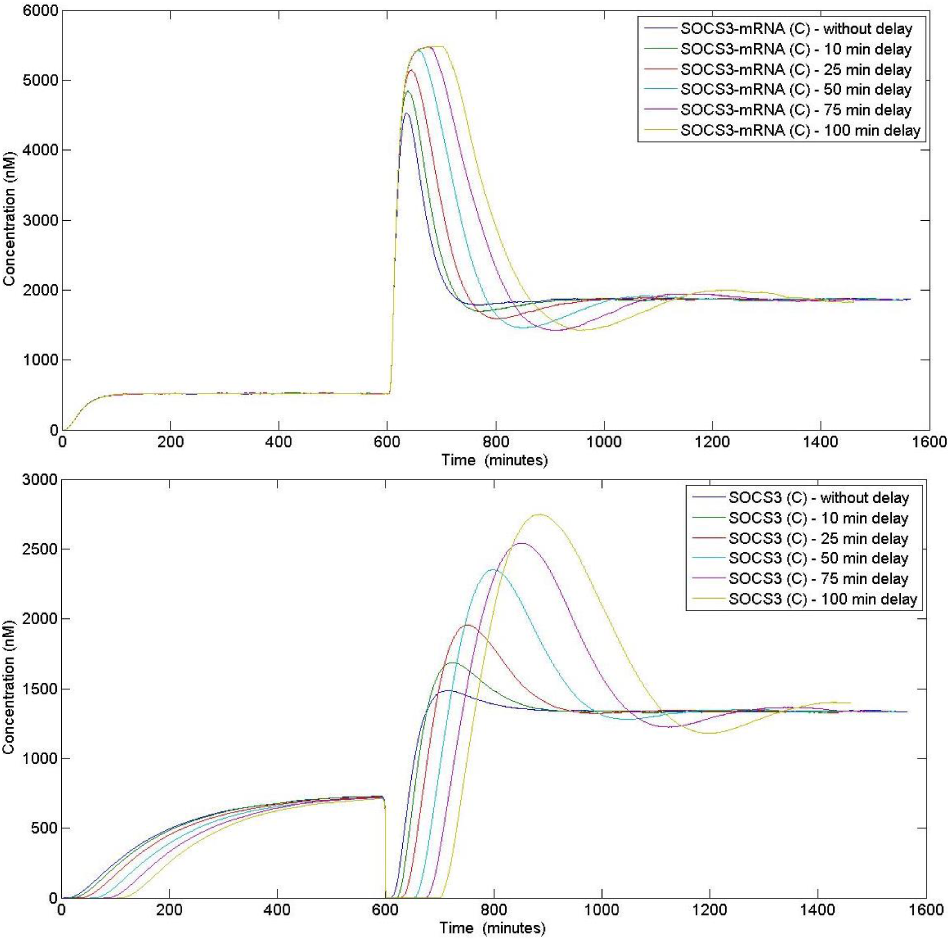
Temporal evolution for SOCS3-mRNA inside the cytoplasm (Up) and for SOCS3 protein inside the cytoplasm (Bottom) for different values of the delay for the translation of SOCS3.

## 5 Conclusion

In this document, we have described the mathematical and computational characteristics of a simple and ready to use implementation of the Gillespie algorithm suitable for simulating the dynamics of stochastic chemical systems with delayed reactions. The implementation and its associated R script is adapted to dynamical systems (re)written as a set of chemical reactions following the law of mass action and assuming that the system is written in terms of elementary reactions only. The practical advantage of this R script is that the source files and especially heavy data structures, such as binary trees, required for this kind of code can be automatically generated. The only assumption is that the model obeys the restrictions on the type of reactions and the order of these reactions.

Although the present code can handle relatively large systems^4^, it remains a serial version only and for simulating very large systems, other implementations (either parallelized or running on GPU for instance) should be used. It is worth emphasizing that what makes stochastic simulations so time consuming is the necessity to simulate hundreds or thousands of times the behavior of the system to obtain correct averaged physical quantities. A straightforward way to speed up the process is to run several batches of runs on different cores rather than using a single core. The performance of this kind of simulator depends on many factors ranging from the kind of machine used to the properties of the system to be simulated such as number of reactions, number of different species and types of reactions. Furthermore, adding delayed reactions can significantly increase the computational time and increasing the total number of molecules within the system decreases the average time step of the algorithm (this is obvious when checking the formula given by eq. 14) which in turn leads to an increase of the computational required for reaching a given physical duration.

Nevertheless, this implementation should be useful for students and researchers familiar with R and Fortran90 and looking for gaining some experience with the use of stochastic simulation algorithms. In all cases, one should keep in mind that when studying the dynamics of biological networks, the simulation itself is usually the very beginning of a long journey during which various kinds of computational and analytical methods might required depending on the purpose of the study and the questions which must be answered.

## Footnotes

1 The term *species* must be understood as holding for any molecular component present, even for a short time, inside the system: a receptor is a species, a ligand is another type of species and the molecule composed by a receptor bound to its ligand is also another species. A given species and its activated form, i.e. after phosphorylation, are considered as two distinct species.

2 As usual each reaction is characterized by a kinetic constant giving the rate of the reaction.

3 For each simulation, it is required to wait until initial equilibrium in absence of LIF is reached. In the corresponding pictures, the timescale has been rescaled accordingly to hide this first part of the dynamic.

4 The present implementation was used to simulate a extended network made of 509 species and 892 elementary chemical reactions obtained by combining different signaling pathways using an extended automatized pipeline

